# Diversity and prevalence of amino acid decarboxylase enzymes in the human gut microbiome – a bioinformatics investigation

**DOI:** 10.1101/2024.12.06.627230

**Authors:** Matthew Sandoval, Dhara D. Shah

## Abstract

Biogenic amines play numerous biological functions that include neuromodulation, maintenance of the gut health and motility, gastric acid secretion, regulation of immune response, cell growth, and gene expression. Therefore, it is crucial to comprehend the potential modulation of these molecules by the human gut microbiota. A primary pathway for the generation of these molecules involves the decarboxylation of amino acids, a process facilitated by enzymes known as amino acid decarboxylases (AADCs). Here, we conducted a bioinformatic analysis to understand diversity and prevalence of AADCs from the most prevalent members of the human gut microbiome. This study aims to understand how human gut microbes generate metabolites that influence health and disease, through specific enzyme activities, with a focus on recognizing the potential role of gut microbiota in neuromodulation, gastrointestinal dysfunctions, immune response regulation, and other critical biological functions. Our results indicate that AADCs are most abundant in the prominent gut microbial genera, namely *Bacteroides*, *Parabacteroides*, *Alistipes*, and *Enterococcus*. Furthermore, among AADCs, arginine decarboxylases are the most common, present in approximately 60% of the frequently found members of the human gut microbiome, followed by aspartate 1-decarboxylases and glutamate decarboxylases. We also found that *Enterococcus faecalis* harbors the most variety of amino acid decarboxylases, potentially playing an important role in driving decarboxylation chemistry in the human gut. In addition, our sequence analyses of various AADCs demonstrated that a tetrad of amino acids in the PLP binding motif can provide functional identification for AADCs. We hypothesize that the diversity in AADCs and the microbes that harbor them has the potential to alter host metabolic outputs. This could provide a mechanism to use specific changes in microbial genera or species to understand possible metabolite modulations that might influence biological functions. Such studies could lay the groundwork for developing future disease markers or therapeutic interventions.

## INTRODUCTION

Amino acid decarboxylases (AADCs) catalyze the decarboxylation of amino acids to generate corresponding amines (Fig 1). Amines produced by these reactions are structurally and functionally diverse and are known as biogenic amines which include trace amines and polyamines. Some AADCs also produce amino acids like GABA. Such molecules play numerous crucial functions in the microbes and in the host harboring these microbes. Specifically, the production of neuromodulatory molecules like histamine, tyramine, tryptamine, dopamine, serotonin, and γ-aminobutyric acid (GABA) is dependent on the actions of AADCs(1–10). Apart from their role in the production of multiple neuromodulatory molecules, AADCs are also involved in the biosynthesis of polyamines like spermine, spermidine, putrescine, and cadaverine(11, 12). Polyamines are produced by the gut microbiota in the large intestine(13, 14). Microbes utilize polyamines for cell growth, during the stress response, and for survival(11). Microbially produced polyamines are also beneficial to the host’s gut health by promoting epithelial renewal, longevity, and recovery of injured mucosa(15, 16). It is also known that in certain gut microbes, the acidification of the gut environment induces polyamine biosynthesis as a coping method for the acidic stress(17). Apart from polyamines, GABA and agmatine also combat acidic stress in bacteria(18, 19). In fact, glutamate and arginine decarboxylases are known to play a role in acid resistance mechanisms present in many prokaryotic organisms(18, 19). Specifically, AADCs consume protons during decarboxylation, increasing the pH and preventing acidic damage to the organism. In addition, polyamines induce glutamate decarboxylase dependent acid resistance systems (20).

**Fig 1.**
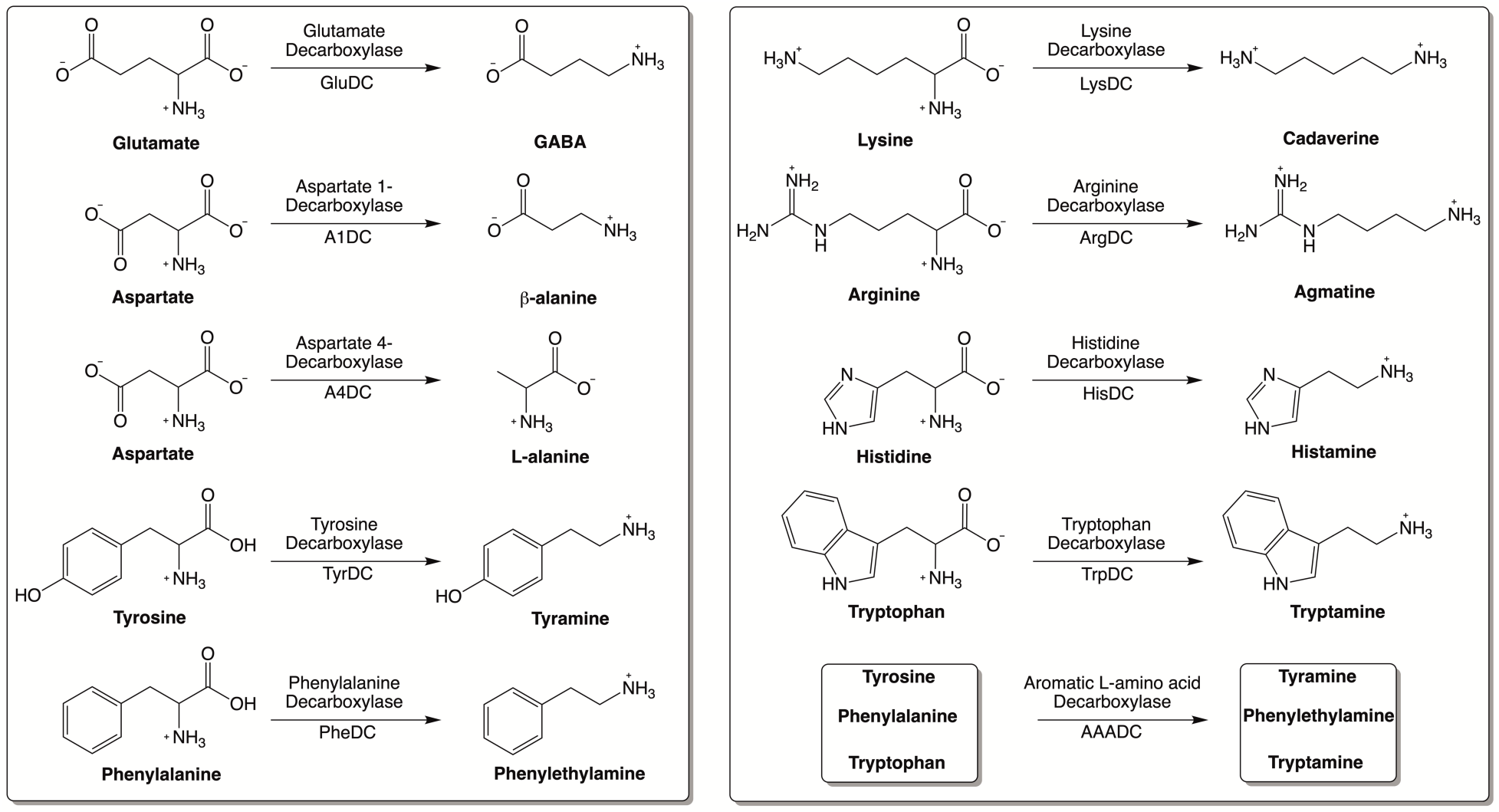
Reaction catalyzed by various amino acid decarboxylases (AADCs).

Moreover, during catalysis by AADCs, carboxylic acid groups are removed from amino acids and released as either CO_2_ gas or as dissolved CO_2_. Depending on the form of the CO_2_ released during the reaction, a variety of physiological changes can occur. As part of a bicarbonate buffering system, it can help maintain the pH within the gut or in the blood stream. In contrast, excessive production of CO_2_ gas can cause discomfort and bloating in humans. For this reason, understanding the nature and abundance of these enzymes in the human gut microbiome can provide useful information about microbial metabolism, communication within microbial communities, and host-microbe interactions. So far, there are numerous isolated studies available on different AADCs which have been critical in comprehending the biochemistry of these reactions(1, 8, 10). In one such study by Williams et al., presence of tryptophan decarboxylases in the human gut microbiota that can generate a neuromodulatory arylamine known as tryptamine was demonstrated(8). Additional studies showed production of other neuromodulatory amines and amino acids like serotonin, tyramine, and GABA generated by the actions of various AADCs present in the human gut microbiota(1, 9, 10, 21). However, the abundance and diversity of various AADCs, the common gut microbes that harbor them, and the collective role of these enzymes in driving decarboxylation chemistry and potentially influencing gut and host physiology, have not been thoroughly investigated or clearly understood. Our systematic study about the prevalence and variety of AADCs of the human gut microbiome provides novel insight into the landscape of decarboxylation chemistry driven by abundant gut microbes harboring these enzymes which might facilitate potential avenues to guide future therapeutic interventions.

## RESULTS AND DISCUSSION

### Arginine decarboxylases are the most abundant amino acid decarboxylases (AADCs) in the prevalent members of the human gut microbiome

Our bioinformatics analysis revealed that arginine decarboxylases were the most represented amino acid decarboxylases among human gut microbiota (Fig 2 and S1 Table). Around 60% of the commonly found gut bacteria harbor arginine decarboxylases (ArgDCs) followed by aspartate-1 decarboxylases (A1DCs) that were present in approximately 50% of the prevalent human gut bacteria (Fig 2). Based on the annotations, these AADCs are predicted to decarboxylate L-arginine and L-aspartate to produce agmatine and β-alanine (Fig 1), respectively. Agmatine is regulator of polyamine biosynthesis and is a precursor for polyamines like putrescine, spermidine and spermine(12). Gut microbes have a metabolic pathway that includes a conserved arginine decarboxylase and a set of other enzymes for the formation of the most abundant polyamine in the gut, spermidine from agmatine(22). Arginine decarboxylases (ArgDCs) play a unique role in gut microbes. They supply agmatine for the production of polyamines in the intestine. In addition, ArgDCs play a crucial role in acid resistance and thus help microbes in the survival under extreme acidic conditions. They facilitate protection by utilizing protons during catalysis and increasing the pH of the solution(7). A1DCs, on the other hand, are important in the formation of β-alanine. β-alanine is a precursor for the biosynthesis of pantothenate that is utilized in the formation of coenzyme A (CoA), an important intermediate in various metabolic pathways(23). β-alanine has also been shown to provide protective effects in individuals with cognitive deficits(24, 25).

**Fig 2.**
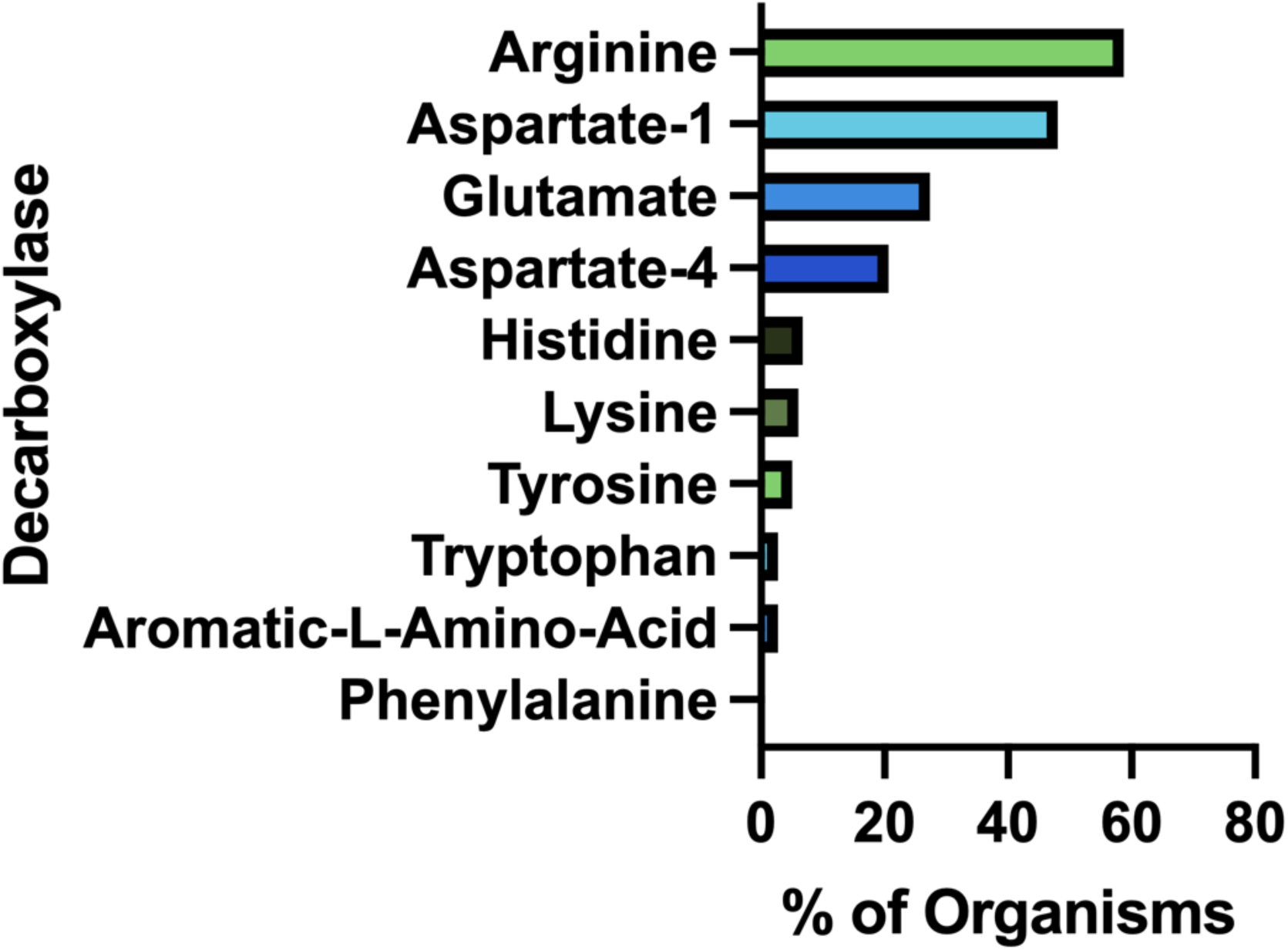
Percent of prevalent human gut microbiota with each type of amino acid decarboxylase (AADC).

The next prominent class of AADCs were glutamate decarboxylases (GluDCs), found to be present in around 27% of the commonly found gut bacteria (Fig 2). Glutamate decarboxylases are known to play an important role in acid resistance mechanism similar to arginine decarboxylases(26). GluDCs catalyze conversion of L-glutamate to γ-aminobutyric acid (GABA) (Fig 1). GABA is the major inhibitory neurotransmitter in the human central nervous system(27). Whereas microbes utilize GABA as an energy source and certain gut microbes grow solely in the presence of GABA(3). With these various functions, glutamate decarboxylases are important AADCs present in the gut microbes. We hypothesize that such overlapping acid resistance mechanisms provided by AADCs like ArgDCs and GluDCs possibly work together in gut microbes. The ability to decarboxylate different amino acid substrates might allow these microbes to survive in various acidic and substrate limiting environments. This strategy allows for continuous acid resistance in microbes specially when one of the pathways become less functional. Following GluDCs in abundance are aspartate-4 decarboxylases (A4DCs). 21% of the gut bacteria contained asparatate-4 decarboxylases (Fig 2). A4DCs catalyze conversion of L-aspartate to L-alanine (Fig 1) and hence A4DCs are important in the metabolism of these two amino acids(4). Moreover, we found that histidine (HisDCs) and lysine decarboxylases (LysDCs) were present in around 7% and 6% of the prevalent human gut bacteria respectively (Fig 2). Histidine decarboxylases (HisDCs) catalyze a conversion of L-histidine to histamine whereas lysine decarboxylases (LysDCs) catalyze a conversion of L-lysine to cadaverine (Fig 1). Histamine plays an important role in communication of immune responses and as a neuroimmune modulator in the gut(28). In contrast, cadaverine produced by the gut microbes can be either detrimental or beneficial to the host(29, 30). Additionally, for some microbes, cadaverine seems to decrease their susceptibility to certain antibiotics(6).

Next to HisDCs and LysDCs, 5% of the prevalent human gut bacteria contained tyrosine decarboxylases (Fig 2). Tyrosine decarboxylases (TyrDCs) catalyze conversion of L-tyrosine to tyramine (Fig 1). In addition, TyrDCs can also covert L-DOPA to dopamine(9, 10). Tyramine is a trace amine and is known to displace catecholamine neurotransmitters like dopamine, epinephrine and norepinephrine from pre-synaptic vesicles and interferes with their signaling(31). Dopamine is a neurotransmitter which is important for motivation, reward, cognition, and motor control(32). Lastly, tryptophan decarboxylases (TrpDCs) and aromatic L-amino acid decarboxylases (AAADCs) were equally represented and found in around 3% of the commonly found human gut bacteria (Fig 2). TrpDCs catalyze conversion of L-tryptophan to tryptamine whereas AAADCs catalyze conversion of aromatic amino acids (tyrosine, tryptophan and phenylalanine) to their corresponding aromatic amines (Fig 1). Tryptamine acts as a neuromodulator in mammalian brain and serves as a regulator of gastrointestinal motility(8). A derivative of tryptamine, 5-hydroxy tryptamine commonly known as serotonin is also an important neurotransmitter(33). AAADCs show broad substrate specificity and other than catalyzing the decarboxylations of three proteinogenic aromatic amino acids, these are known to also decarboxylate derivatives of aromatic amino acids like 3,4-dihydroxyphenylalanine (L-DOPA)(9, 10) and 5-hydroxytryptophan to produce dopamine and 5-hydroxytryptamine (serotonin)(34). We did not find separately annotated phenylalanine decarboxylases (PheDCs) in any of the common gut microbes. We hypothesize that if there are enzymes which decarboxylate phenylalanine, then these are found under the bigger class of enzymes called aromatic L-amino acid decarboxylases(1). The decarboxylation product of L-phenylalanine is phenylethylamine (PEA) (Fig 1). PEA like many other trace amines can bind to trace amine-associated receptor 1 and impart various physiological effects(35, 36). Some of these are, activation of blood leukocytes(35) and alteration of monoamine transporter function in the brain(36). Given the diverse biological roles of the products generated by AADC-mediated reactions, gaining insight into their distribution among human gut microbes may inform future methods for modulating concentrations of these compounds in humans.

### The prevalent gut microbial genus *Bacteroides* has the highest abundance of amino acid decarboxylases (AADCs)

In our investigation of annotated amino acid decarboxylases among common human gut bacteria, we observed a broad spectrum of abundance levels for various classes of amino acid decarboxylases (AADCs) within each genus, as shown in Fig 3. The detected numbers varied widely, from as many as 60 to as few as none. Our analysis showed that the genus *Bacteroides* contained the highest number of AADCs (Fig 3). We found 63 annotated AADCs in the members of *Bacteroides* genus which were significantly higher than any other gut microbial genus (Fig 3 and S2 Table). The next 5 genus after *Bacteroides* showed anywhere between 9-18 AADCs. These were *Enterococcus* with 18, *Alistipes* with 14, *Parabacteroides* with 12, *Streptococcus* with 10, and *Prevotella* with 9 AADCs (Fig 3 and S2 Table). All the top 6 genera belong to two phyla Bacteroidetes and Firmicutes. Interestingly, these are also the two major phyla that produce CO_2_ and constitute over 90% of the total microbial population of the human gut(37). The next three genera *Enterobacter* (Proteobacteria), *Klebsiella* (Proteobacteria), and *Phocaeicola* (Bacteroidetes) each contain total 7 AADCs (Fig 3 and S2 Table). There were 5 genera that showed the presence of 6 AADCs, 7 genera that had 4 AADCs, and 15 genera that had 3 AADCs as highlighted in Fig 3. Apart from that, 48 genera contained either 1 or 2 AADCs and 5 genera did not have any AADCs (S2 Table). *Bacteroides* represents one of the most prevalent genera within the human gut microbiome, leading to a higher number of its species and strains being well characterized relative to other genera. Within the gut microbes examined in this study, multiple *Bacteroides* species are found in significant numbers. Therefore, the results presented here may be influenced by the extensive species data from *Bacteroides*, as compared to data from other microbial genera in the human gut.

**Fig 3.**
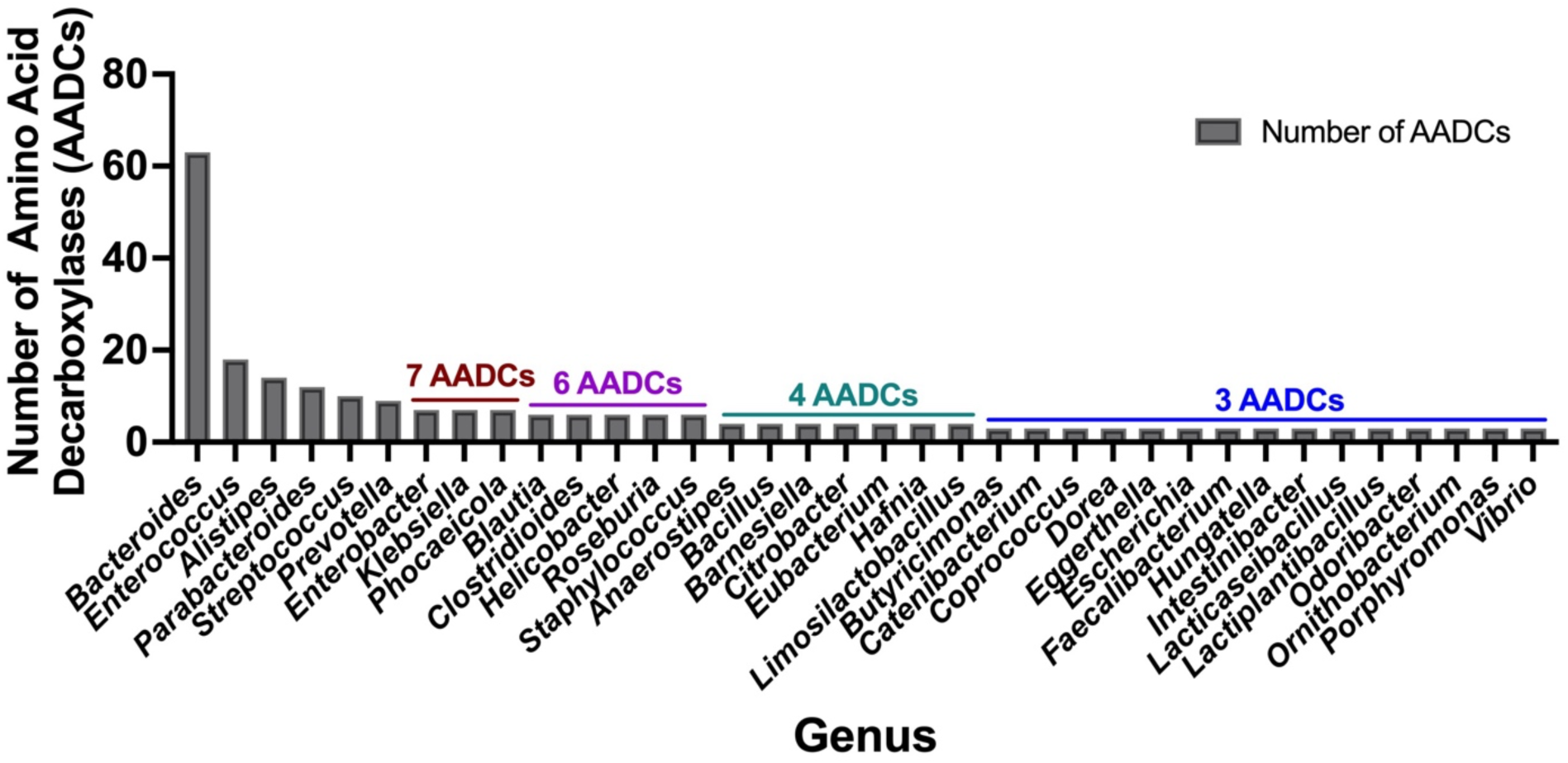
Total number of amino acid decarboxylases (AADCs) at genus level. The plot depicts genus level occurrences of all classes of amino acid decarboxylases in the prevalent human gut bacteria. Only genera harboring ≥ 3 AADCs are included here. The entire list for genus level AADCs occurrences are presented in S2 Table.

### *Enterococcus faecalis* harbors the most variety of amino acid decarboxylases

Next, we performed an analysis to understand if there are any patterns in the type of AADCs present in the prominent gut microbes. Our results demonstrated that the type of AADCs present were not genus specific and different species within the same genus can contain different classes of AADCs (Fig 4 and S3 Table). We found that *Enterococcus faecalis* had the most different classes of AADCs. It contained 7 out of 10 AADCs analyzed in this study. However, the genus *Enterococcus* represented anywhere from 1 to 7 types of AADCs at the species level (Fig 4 and S3 Table).

**Fig 4.**
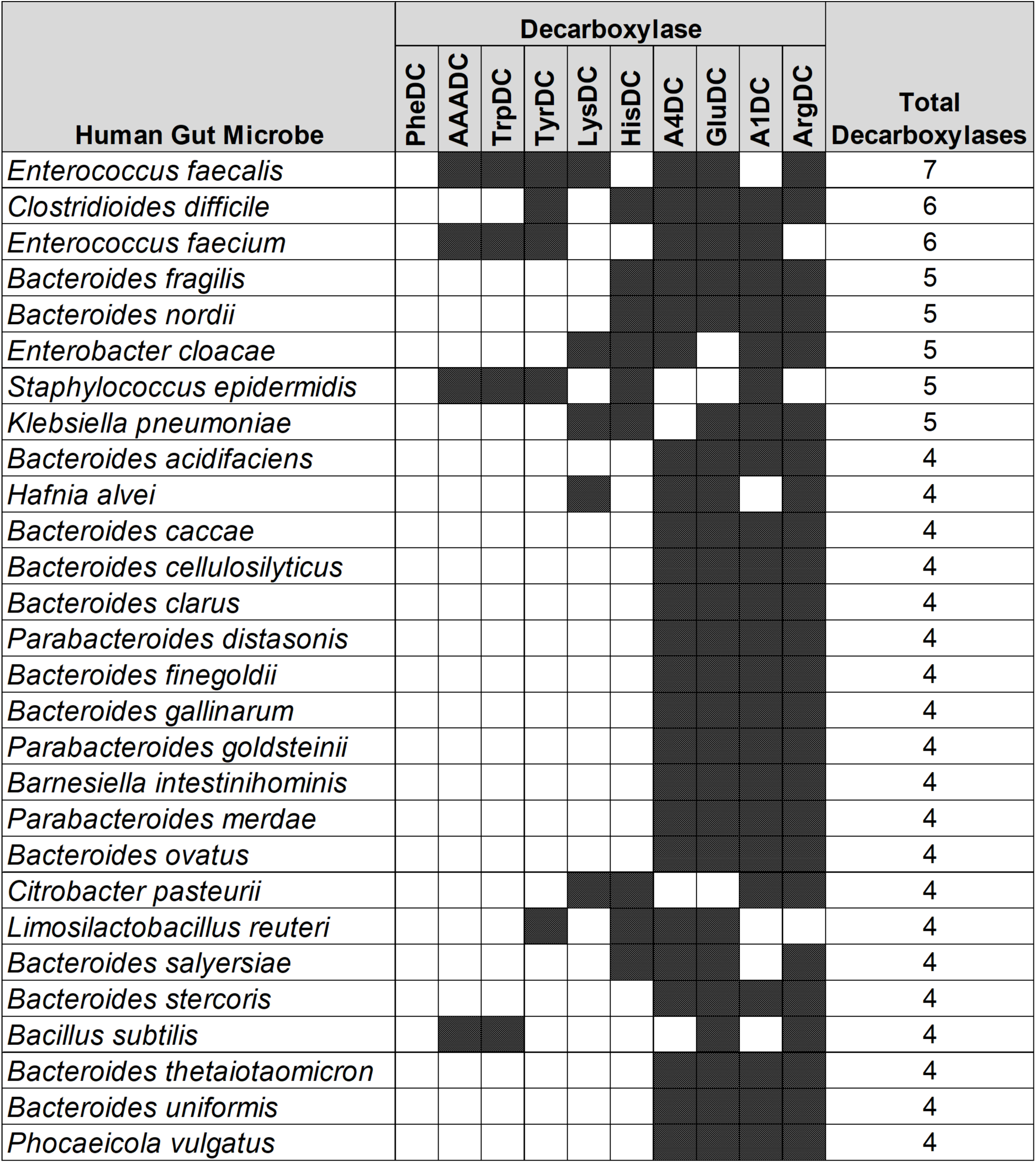
Different classes of AADCs present in the most commonly found human gut microbes. The figure represents prevalent human gut microbes harboring different classes of AADCs. Here, shaded areas correspond to the presence of AADCs, and non-shaded (blank white) areas correspond to the absence of AADCs. Only microbes containing at least 4 different classes of AADCs have been included. The full list of the commonly found gut microbes containing various classes of AADCs can be found in S3 Table.

A similar variation, but not as large, was also seen with the genus *Bacteroides*. We found that the various species of *Bacteroides* contained 3 to 5 different classes of AADCs where the majority of the species had 4 different types of AADCs (Fig 5). It is interesting to note that the genus *Bacteroides* showed the presence of at least 3 classes of AADCs and this points towards the importance of AADCs in their metabolism. Fig 5 depicts the presence of different AADCs in various species of the genus *Bacteroides*. All the species of *Bacteroides* present in this study have GluDC, ArgDC, and A4DC. The other remaining classes of AADCs found in some of the species are A1DCs and HisDCs. HisDCs are the least common in Bacteroides and only present in three *Bacteroides* species. Additionally, we observed that the genus *Bacteroides* lack AADCs that are able to catalyze decarboxylations of aromatic amino acids – tyrosine, tryptophan, and phenylalanine. These reactions are generally catalyzed by TyrDC, TrpDC, PheDC, and AAADC. This points towards the inability of the genus *Bacteroides* to produce aromatic amines specifically via the route of decarboxylations of aromatic amino acids. In addition to AAADCs, *Bacteroides* also lack lysine decarboxylase (LysDCs).

**Fig 5.**
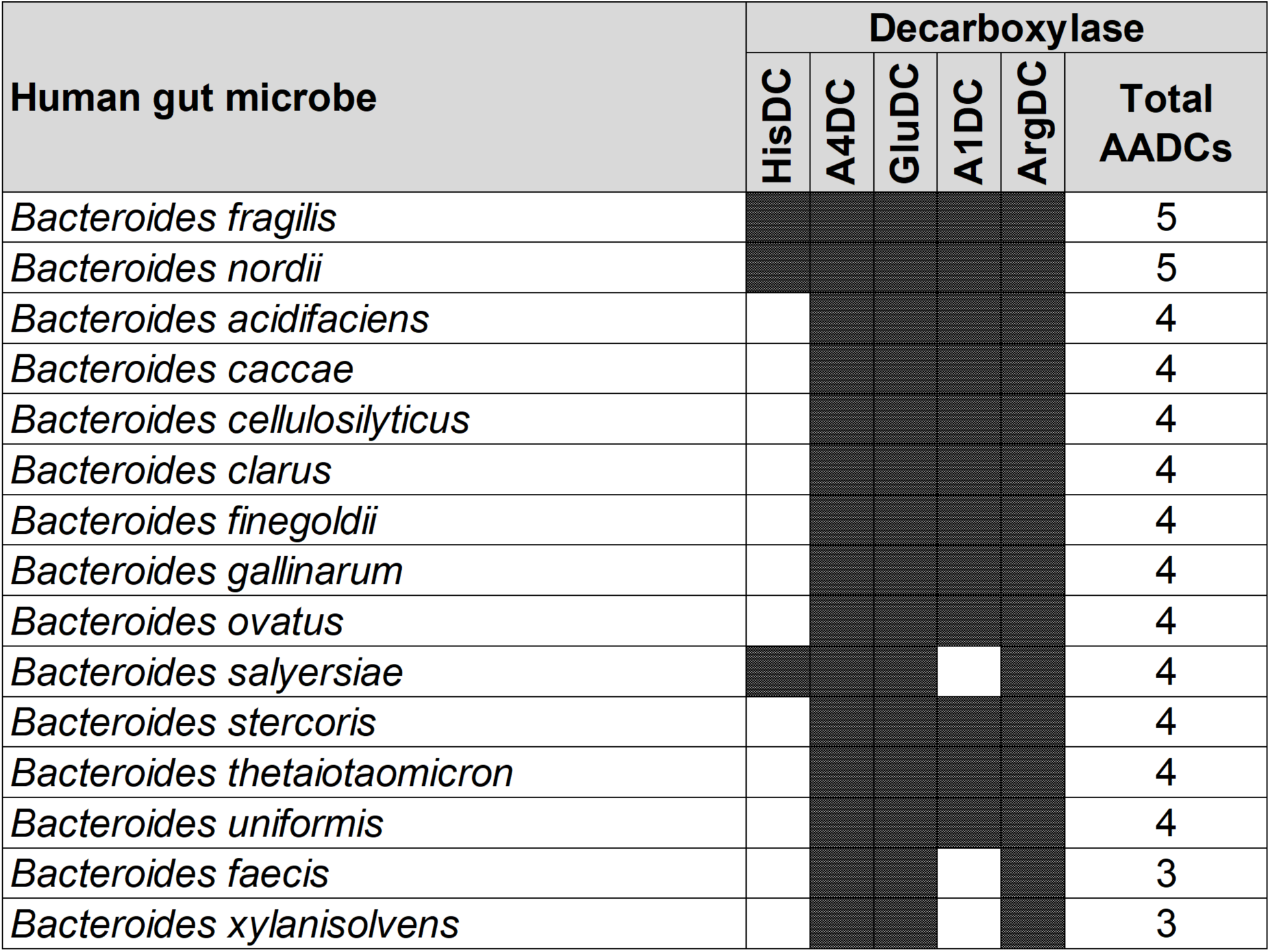
Different classes of AADCs present in various species of the genus *Bacteroides.* The figure depicts the presence of different classes of AADCs in various species of the genus Bacteroides. The shaded areas in the figure show the presence of AADCs, and non-shaded (blank white) areas show the absence of AADCs.

S3 Table represents a full list of prevalent human gut microbes with or without the presence of 10 classes of AADCs. In addition to microbes harboring 4 or more AADCs depicted in Fig 4, there were 50 bacteria containing at least one type of AADC, 40 bacteria with two different types of AADCs, and 20 bacteria with 3 different types of AADCs (S3 Table). Moreover, we found that the genus *Bifidobacterium* severely lacked AADCs. Other than the one species – *B. adolescentis* with one AADC which is GluDC(2), none other members of the genus had any of the 10 AADCs. During our analysis, we observed variability at the strain level in a few instances among identical species of the microbes. The scope of this study is beyond the strain level variation. However, our results indicated that the types of AADCs can vary at the genus, species, and/or strain level. This variability in AADCs can potentially affect microbial metabolism, specifically the ability to generate biogenic amines including trace amines, polyamines and, amino acids like GABA with varying biological functions either towards the community of other gut microbes or towards the host. In addition, this variability might drive variations in the production of carbon dioxide that can impart either advantageous or detrimental effects to the host some of which are impacting pH balance, changes in gut motility, and gas accumulation and bloating.

### Arginine decarboxylases (ArgDCs) display the most significant differences in length among the human gut microbes

To understand if the AADCs found in various gut microbes are similar or different, we conducted a protein length analysis on all the AADCs found in the common human gut microbes (S4 Table). Through this analysis, we found that TyrDCs and A1DCs shared similar length across different genera and species (Table 1). These were followed by HisDCs and GluDCs. AADCs from these two classes show variability of around 80 to 90 residues in the protein length across prevalent human gut microbial genera and species. However, A4DCs, AAADCs, and TrpDCs showed larger variation in the length which was from 120 to 180 residues. Our dataset with AAADCs and TrpDCs is very small that contains total five enzymes only. Interestingly, AAADCs and TrpDCs were exactly same which points towards two possible annotations provided as AAADCs and TrpDCs. LysDCs showed even greater range in protein lengths which was around 300 residues that contained enzyme with a maximum length of around 740 amino acids and the minimum length of around 450 amino acids (Table 1). The last one and with the biggest range in the length is represented by ArgDCs. The protein length range was found to be around 640 residues that contained the longest ArgDC with 790 amino acid residues and the shortest ArgDC with 154 amino acid residues (Table 1). The majority of proteins in ArgDC family had around 630 (mode value) amino acid residues. There are four known classes of prokaryotic arginine decarboxylases (ArgDCs)(7). We think that the variability in protein lengths is due to the presence of various types of ArgDCs within human gut microbes. Specifically, the type of ArgDCs present in Bacteroidetes and Firmicutes are different from each other. Bacteroidetes have alanine racemase fold ArgDC (AR-fold ArgDC) whereas firmicutes have aspartate amino transferase fold ArgDC (AAT-fold ArgDC)(7). Additionally, ArgDCs have different structures based on their biological functions. Acid inducible ArgDCs are important during acid stress in microbes whereas the other class of ArgDCs are not upregulated during acid stress and their main biological function is in polyamine biosynthesis(7).

**Table 1.** Length variation within amino acid decarboxylases.

| AADC | Enzyme ID | Length Range | Length Max | Length Min | Mode | Number of Enzymes |
| --- | --- | --- | --- | --- | --- | --- |
| ArgDC | 4.1.1.19 | 636 | 790 | 154 | 630 | 107 |
| LysDC | 4.1.1.18 | 289 | 739 | 450 | 712 | 12 |
| TrpDC | 4.1.1.105 | 178 | 513 | 335 | 470-484 | 5 |
| AAADC | 4.1.1.28 | 178 | 513 | 335 | 470-484 | 5 |
| A4DC | 4.1.1.12 | 121 | 588 | 467 | 547 | 38 |
| GluDC | 4.1.1.15 | 92 | 551 | 459 | 480 | 51 |
| HisDC | 4.1.1.22 | 85 | 380 | 295 | 295 | 11 |
| A1DC | 4.1.1.11 | 29 | 142 | 113 | 117 | 87 |
| TyrDC | 4.1.1.25 | 20 | 635 | 615 | 625 | 9 |

Analysis in this table is based on the amino acid sequence data from S4 Table.

### Sequence similarities within AADCs from each class provide either functional significance or the microbial host specificity

Our analysis presents interesting aspects about decarboxylase enzymes within each group of AADCs. As mentioned above, there are multiple types of ArgDCs present in prokaryotic organisms. Previous, studies have already identified that ArgDCs are phyla specific(7). This is reflected in the percent identity matrix for all ArgDCs where a few areas with high sequence similarities are visible (S5 Table). For the other large group GluDCs, we found that there are two major areas with high sequence similarities in the percent identity matrix (S6 Table). The area highlighted with yellow outline mostly contain genera within the phylum Firmicutes (now Bacillota) which contain gram-positive organisms. However, the other area highlighted with green outline mostly represents organisms that belong to the phylum Bacteroidetes (now Bacteroidota) with two exceptions of *Faecalibacterium prausnitzii* and *Catenibacterium mitsuokai*. Based on these results, gut microbial GluDCs from the phylum Bacteroidetes are more similar to each other than to GluDCs of the phylum Firmicutes. When we compare GluDCs of Firmicutes to GluDCs of Bacteroidetes, we still see around 40 – 50% overall sequence similarity which is significant (S6 Table). Unlike ArgDCs where completely different classes of enzymes are present in Firmicutes and Bacteroidetes, here we see more subtle differences which are present in the overall amino acid sequences. Moreover, in the percent identity matrix of GluDCs, we found two enzymes that are significantly different from the whole group with only 15 – 20% overall sequence similarity to the other members of the group. These were from *Pararheinheimera texasensis* and *Vibrio cincinnatiensis*. Our analysis suggests that these might be misannotated and could belong to some other class of AADCs. We saw a similar pattern with A1DCs where we observed three distinct areas with high sequence similarities. These areas are highlighted in S7 Table, where the top left most (highlighted in yellow) contain organisms from the phylum Proteobacteria (now pseudomondota) which are primarily gram negative and facultative anaerobes. The middle area with high sequence similarity (highlighted in green) was occupied mostly by the members of the phylum Firmicutes (now Bacillota) which are gram positive and anaerobic organisms. We did see some exceptions in that area. The last area with the high sequence similarity was the right bottom area (highlighted in cyan) that represented members of the phylum Bacteroidetes (now Bacteroidota), which are gram negative and anaerobic organisms.

A4DCs are not as prevalent as other classes of AADCs discussed above but in the percent identity matrix we see again a region with high sequence similarity that is coming from the members of the phylum Bacteroidetes (highlighted in cyan) (S8 Table). Additionally, two groups of AADCs are exactly same and contain exact same members and proteins. These two are AAADCs and TrpDCs (S9 Table). Interestingly, these enzymes were mostly seen in the members of the phylum Firmicutes (now Bacillota). We found one member of the phylum Actinobacteria (now Actinomycetota), *Nitriliruptor alkaliphilus* harboring AAADC/TrpDC. Of the other smaller groups of AADCs, HisDCs were present equally in both phyla, Bacteroidetes and Firmicutes (S10 Table). The percent identity matrix (S10 Table) displayed two HisDCs which shared very low sequence similarity with the rest of the group. These were present in, *Enterobacter cloacae* and *Klebsiella pneumoniae* which are both Proteobacteria. The overall sequence similarity of these two with the whole group of HisDCs is only around 14 – 20%. We hypothesize that these might be misannotated and could belong to some other class of AADCs. The next AADCs class that is similar in prevalence as HisDCs is LysDCs. We found that LysDCs are mostly present in the members of the phylum Proteobacteria (now Pseudomonadota) with only two members from the phylum Firmicutes (now Bacillota) (S11 Table). In addition, LysDCs showed very high sequence similarity within the group with the average percent identity of around 70% (Table 2). However, a LysDC from *Peptoniphilus harei* was found to be very different from the rest of the group with an overall sequence similarity of only 25 – 27% (S11 Table). Based on this information, there is a possibility that the enzyme was misannotated as HisDC. We think that the enzyme might belong to the other group of AADCs and an experimental validation will be necessary to identify its true function. The last group of AADCs that is smaller in size is TyrDCs. Our analysis revealed that TyrDCs are mostly present in the members of the phylum Firmicutes (now Bacillota) (S12 Table). We found only one enzyme that was from *Cutibacterium acnes* which is an Actinobacteria. Interestingly, TyrDC from this organism showed the lowest sequence similarity to all the other members of the group. However, the overall sequence similarity of this enzyme to all the other members of the TyrDC class is still high, around 42 – 45%. This enzyme represents the sole instance of TyrDC from the Actinobacteria, while all other TyrDCs originate from Firmicutes, and this could be the possible reason behind sequence divergence in TyrDCs. This class exhibited the greatest sequence similarity as a group when contrasted with any other group of AADCs (Table 2).

**Table 2.** Percent identity based on the amino acid sequence similarities within each group of AADCs.

| Amino acid Decarboxylase | Average Percent Identity | Minimum Percent Identity |
| --- | --- | --- |
| Arginine Decarboxylase | N/A | N/A |
| Tryptophan Decarboxylase | 50.86 | 24.84 |
| Histidine Decarboxylase | 44.08 | 30.34 |
| Aromatic-L amino acid Decarboxylase | 50.86 | 24.84 |
| Glutamate Decarboxylase | 55.73 | 36.03 |
| Aspartate-1 Decarboxylase | 58.68 | 31.97 |
| Aspartate-4 Decarboxylase | 57.14 | 35.58 |
| Lysine Decarboxylase | 69.44 | 61.88 |
| Tyrosine Decarboxylase | 72.18 | 42.83 |

### A tetrad of amino acids in the PLP binding motif can provide functional identification and assignment for most amino acid decarboxylases (AADCs)

Next, we explored the extent of variation among these AADCs in the regions surrounding their cofactor binding sites. Most known amino acid decarboxylases studied here utilize PLP as the cofactor to catalyze decarboxylation reactions. PLP cofactor binds to amino acid decarboxylases via a strictly conserved lysine residue and this lysine is present in all PLP dependent decarboxylases(38). Our analysis of the partial PLP binding motifs specifically four amino acids surrounding the conserved lysine residue showed that within each group of AADCs, these tetrads are conserved (Table 3). Our examination of the partial PLP binding motifs, focusing on the four amino acids around the conserved lysine, revealed that these tetrads are preserved within each AADC group. For GluDCs, we see a conserved amino acid tetrad of “SGHK” (Table 3). Based on our percent identity matrices, we saw that there were two outliers in GluDCs group, *Pararheinheimera texasensis* and *Vibrio cincinnatiensis*. The PLP binding motif analysis for these two is in agreement with the sequence similarity analysis (S13 Table). We observed that the two outliers had a different PLP conserved tetrad of “DAHK”. This strengthens our hypothesis that the AADCs from *Pararheinheimera texasensis* and *Vibrio cincinnatiensis* are possibly not GluDCs. While looking through the PLP motifs of other AADCs, we found that DAHK motif is generally present in AAADCs/TrpDCs family (Table 3 and S14 Table). Due to the presence of this motif in AADCs of *P*. *texasensis and V. cincinnatiensis,* these might possibly be aromatic L-amino acid decarboxylases. In addition to our analysis, experimental evidence collected by *in vitro* characterization of purified enzymes can provide correct functions of these AADCs. For the aromatic-L-amino acid decarboxylases group that also contains tryptophan decarboxylases, the conserved tetrad residues with variations found at the first and the second positions were “[DN]-[AP]-HK”, where we find motifs like DAHK, DPHK, NPHK, and NAHK (Table 3 and S14 Table). These variations might be useful in understanding specificity of AAADCs towards specific aromatic amino acids. One such example is in the Table 3, where we see a conserved tetrad of “DPHK” for TyrDCs. We did not find any exceptions in this conserved motif in annotated TyrDCs (S15 Table). It is possible that decarboxylases under the group AAADCs harboring DPHK motifs might be TyrDCs. However, this will need a thorough experimental investigation. For A4DCs, we found conserved tetrad residues as “S-[Fy]-[Sa]-K” with variations found at the first and the second positions (Table 3). The primary and most commonly found motif was “SFSK”, followed by “SYSK” found in only 5.3 % of enzymes of this class and lastly the motif “SFAK” found in only 2.6 % enzymes of this class (Table 3 and S16 Table). The next group of enzymes harboring conserved tetrad was LysDCs. We found a conserved tetrad of “N-[Ct]-HK” with variations observed at the second position (Table 3). Here, the most found motif was “NCHK”. In only one instance, we found the alternate motif of “NTHK” (S17 Table). However, this motif was present in an enzyme from *Peptoniphilus harei* that was an outlier during our sequence similarity analysis. The motif is also present at a different position (residues 99-102) compared to all the other enzymes in the group which showed motif within amino acid residues 243-246 (254-257 in one case). These observations point towards the possibility of the enzyme from *P. harei* to be AADC from another group or an unusual LysDC. Experimental evidence is required in understanding the true function and nature of this enzyme. All identifiable PLP binding motifs shared the essential lysine (K) residue in similar positions. This residue was most preceded by a histidine (H) residue but was replaced with mostly serine (S) and rarely with alanine (A) resides within the group asparate-4 decarboxylases (A4DCs).

**Table 3.**
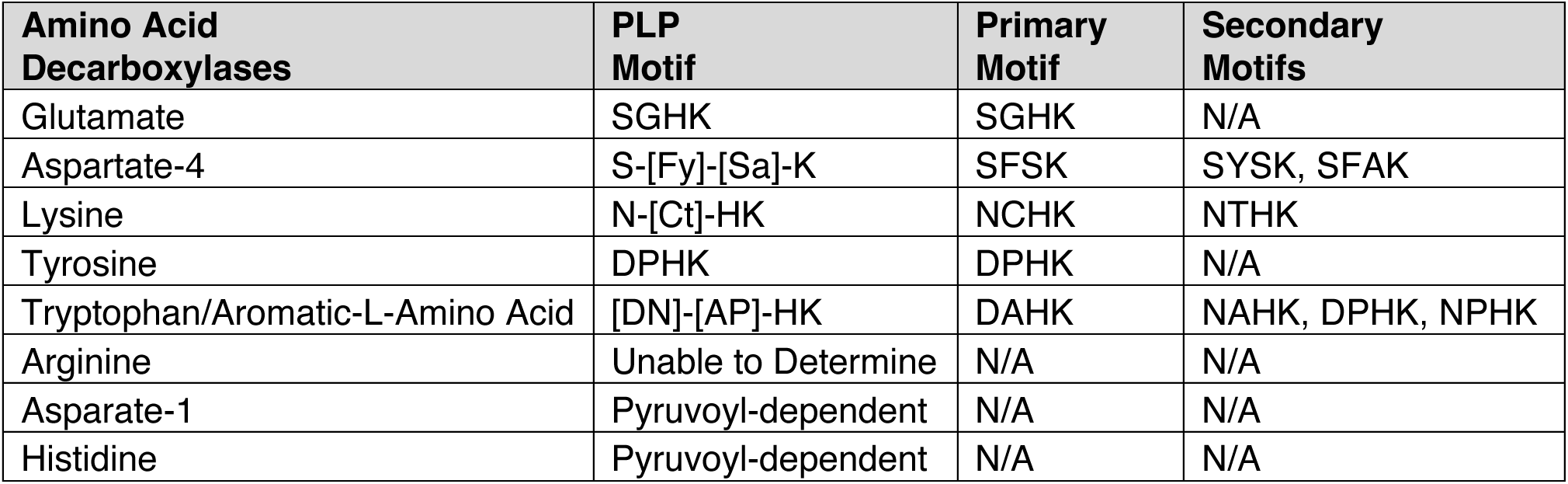
PLP binding motif in AADCs of common human gut microbiota.

Apart from the PLP dependent amino acid decarboxylases, another class of AADCs are also present in prokaryotes, known as pyruvoyl-dependent amino acid decarboxylases. There are three known AADC groups that contain pyruvoyl cofactor in place of PLP cofactor. These are histidine decarboxylases (HisDCs)(39–42), aspartate 1 decarboxylases (A1DCs)(43) and some arginine decarboxylases (ArgDCs)(44). ArgDCs have been classified previously in many different groups and due to this variation, we were unable to find conserved PLP motifs in ArgDCs(7). Similarly, we only found pyruvoyl dependent HisDCs and A1DCs in the prevalent members of the human gut microbiome and hence no PLP motif was found within these enzymes. Interestingly, human histidine decarboxylase is a PLP dependent enzyme(45) unlike the enzymes of the human gut microbes which are pyruvoyl-dependent(46). While both types of enzymes produce histamine, the underlying mechanisms differ between human enzymes and those found in gut microbes, which could result in variations in activity and regulatory control. In contrast to HisDCs, humans do not have an aspartate decarboxylase (A1DC) that produces β-alanine unlike the members of the human gut microbiome. Humans produce β-alanine via a separate metabolic pathway and not via the direct decarboxylation of L-aspartate(47). β-alanine is the precursor for the dipeptide carnosine which is found in muscles and can help combat muscular fatigue during strenuous exercise(48). Additionally, β-alanine has been identified as a neurotransmitter and is an essential component of the coenzyme A(47, 49). However, recently an enzyme called GADL1 (glutamic acid decarboxylase like 1) showed a direct decarboxylation activity with L-aspartate to produce β-alanine(48). This enzyme is a PLP dependent enzyme in contrast to pyruvoyl dependent gut microbial enzymes.

## CONCLUSION

Our *in-silico* analysis of gut microbial AADCs provides a useful understanding about the prevalence, conservation, functional context, and host specificity of these enzymes. With the study presented here, we also show the need to understand the role of these decarboxylases in-depth. The research in this field can provide crucial information about the functional significance of AADCs for each microbe harboring these enzymes, possible roles of these enzymes including communication within the gut microbial community, and ultimately the possible roles of AADCs in host-microbe interactions and modulations of important metabolites that affect human metabolism and health. However, this study does not underscore the necessity to understand the enzymatic activities *in-vitro* and *in-vivo*, which will be crucial to study AADCs within their natural context.

## METHODS

### Database

This research was conducted using the Integrated Microbial Genomes & Microbiomes (IMG/M) public facing database for genome datasets(50).

### Selection of gut microbial candidates

Human gut bacterial candidates were selected from a previously published human gut bacterial genome and culture collection studies by Forster et al.(51). This study identified human gut microbiota through fecal samples collected from 20 healthy adults from North America and the United Kingdom who had not recently taken antibiotics. Species selection for investigating amino acid decarboxylases was conducted using this data and included human gut bacteria that were present at levels greater than 0.01% within any two samples analyzed as presented in the Supplementary Table 5 from the study conducted by Forster et al.(51). Species not identified as the member of common human gut microbiota by Forster et al. were not selected. After species selection, each species was inquired by Taxon ID through IMG/M and alternative species names (recently changed names) were identified and recorded (S18 Table).

### Enzyme ID (EC number) selection

The IGM/M database was inquired for “decarboxylase” and all amino acid decarboxylases of interest were identified and enzyme IDs were saved (S19 Table). No exclusion criteria were applied at this stage.

### Extraction of gene ID data harboring amino acid decarboxylases

Each EC number (Enzyme ID) was queried in IMG/M and all gene IDs harboring a specific EC number were exported. All bacteria with an annotated amino acid decarboxylase of interest were included and no exclusion criteria were applied. All bacteria (present in the IMG/M database) containing various amino acid decarboxylases were grouped together under a specific Enzyme ID (S1 File).

### Refinement of gene ID data with the available abundant human gut microbes

The gene ID data for each EC number (Enzyme ID) was filtered for only those that corresponded to a common human gut bacterial species (or alternative species name as outlined in species selection) selected from the Forster et al.(51) data as mentioned above. Given the quantity of gene IDs from each species due to the presence of multiple strains, a representative gene ID was selected from the collection with preference for complete genomes present in either ATCC, NCTC, or DSM culture collections. When multiple annotations for a species’ decarboxylase were present in ATCC, NCTC, or DSM, the representative was selected without preference. For species without an annotated genome in ATCC, NCTC, or DSM, an alternative gene ID was selected as the representative without preference (S2 File).

### Representative species selection

Within each decarboxylase group, one species was selected to have all strains (with different Gene IDs) harboring the decarboxylase enzyme evaluated (highlighted in green under each Enzyme ID in the (S2 File). This was necessary to confirm that the selection process for a representative strain for each species without preference, outside of ATCC, NCTC, or DSM annotations, was adequate to become representative for strains within one species. Amino acid sequence homology within the decarboxylase enzyme from various strains of the same species were very high, well above 90%, in almost all cases (S20 Table). This homology between strains of the same species indicates that the selection of one strain per species was an appropriate method for identifying decarboxylase sequences to be analyzed. There was only one case where the sequence homology was poor. It was found within the tryptophan decarboxylases of *N. alkaliphilus* strains which might be indicative of greater variation within *N. alkaliphilus* enzymes. Specially because annotations for these enzymes are either aromatic amino acid decarboxylase or glutamate or tyrosine decarboxylase. In such cases, without the biochemical characterization of these enzymes, discerning functions will not be possible.

### Multiple sequence alignment

After the identification of a gene ID for each species, the amino acid sequence data for the protein product of each gene ID was exported from IMG/M and saved for further analysis in S3 File. All amino acid decarboxylase sequences were obtained from IMG/M. The sequence data for each decarboxylase was saved as a FASTA file with the gene ID and species name in the header. Multiple Sequence Alignment was then performed on each group of decarboxylases using ClustalW Omega 1.2.4 with default parameters to evaluate homology of the various human gut microbial amino acid decarboxylases. Alignment outputs were saved, and the Percent Identity Matrix (PIM) file was exported and formatted in Microsoft Excel to visualize similarity between decarboxylases across genera and species.

### Identification of PLP (pyridoxal phosphate) binding motifs within each amino acid decarboxylase group

The PLP Binding motif was identified through the common motif found in many PLP dependent decarboxylases(38) from the multiple sequence alignment data. The motif was uniquely identified by its position within the peptide sequence and a characteristic Lysine (K) residue (S13 Table – S17 Table).

## SUPPORTING INFORMATION

**S1 Table. Percent of microbes with each type of AADC.**

**S2 Table. Genus level occurrences of AADCs.**

**S3 Table. Types of AADCs in gut microbes.**

**S4 Table. amino acid sequences.**

**S5 Table. Percent identity ArgDCs.**

**S6 Table. Percent identity GluDCs.**

**S7 Table. Percent identity A1DCs.**

**S8 Table. Percent identity A4DCs.**

**S9 Table. Percent identity AAADCs and TrpDCs.**

**S10 Table. Percent identity HisDCs.**

**S11 Table. Percent identity LysDCs.**

**S12 Table. Percent identity TyrDCs.**

**S13 Table. PLP motif GluDCs.**

**S14 Table. PLP motif AAADCs&TrpDCs.**

**S15 Table. PLP motif TyrDCs.**

**S16 Table. PLP motif A4DCs.**

**S17 Table. PLP motif LysDCs.**

**S19 Table. Decarboxylase enzymes.**

**S20 Table. Percent identity in amino acid sequences of each type of decarboxylase from the different strains of the same species.**

**S1 File. EC ID Search Raw Data.**

**S2 File. EC ID Search Gene ID Data.**

**S3 File. Amino acid sequences for decarboxylases.**

## Notes

### Competing Interest Statement

The authors have declared no competing interest.

## REFERENCES

1. Sugiyama Y, Mori Y, Nara M, Kotani Y, Nagai E, Kawada H, et al. Gut bacterial aromatic amine production: aromatic amino acid decarboxylase and its effects on peripheral serotonin production. Gut Microbes. 2022;14(1):2128605.

2. Duranti S, Ruiz L, Lugli GA, Tames H, Milani C, Mancabelli L, et al. Bifidobacterium adolescentis as a key member of the human gut microbiota in the production of GABA. Sci Rep. 2020;10(1):14112.

3. Strandwitz P, Kim KH, Terekhova D, Liu JK, Sharma A, Levering J, et al. GABA-modulating bacteria of the human gut microbiota. Nat Microbiol. 2019;4(3):396–403.

4. Wang NC, Lee CY. Enhanced transaminase activity of a bifunctional L-aspartate 4-decarboxylase. Biochem Biophys Res Commun. 2007;356(2):368–73.

5. Wuthrich D, Berthoud H, Wechsler D, Eugster E, Irmler S, Bruggmann R. The Histidine Decarboxylase Gene Cluster of Lactobacillus parabuchneri Was Gained by Horizontal Gene Transfer and Is Mobile within the Species. Front Microbiol. 2017;8:218.

6. Akhova A, Nesterova L, Shumkov M, Tkachenko A. Cadaverine biosynthesis contributes to decreased Escherichia coli susceptibility to antibiotics. Res Microbiol. 2021;172(7-8):103881.

7. Burrell M, Hanfrey CC, Murray EJ, Stanley-Wall NR, Michael AJ. Evolution and multiplicity of arginine decarboxylases in polyamine biosynthesis and essential role in Bacillus subtilis biofilm formation. J Biol Chem. 2010;285(50):39224–38.

8. Williams BB, Van Benschoten AH, Cimermancic P, Donia MS, Zimmermann M, Taketani M, et al. Discovery and characterization of gut microbiota decarboxylases that can produce the neurotransmitter tryptamine. Cell Host Microbe. 2014;16(4):495–503.

9. Maini Rekdal V, Bess EN, Bisanz JE, Turnbaugh PJ, Balskus EP. Discovery and inhibition of an interspecies gut bacterial pathway for Levodopa metabolism. Science. 2019;364(6445).

10. van Kessel SP, Frye AK, El-Gendy AO, Castejon M, Keshavarzian A, van Dijk G, et al. Gut bacterial tyrosine decarboxylases restrict levels of levodopa in the treatment of Parkinson’s disease. Nat Commun. 2019;10(1):310.

11. Gevrekci AO. The roles of polyamines in microorganisms. World J Microbiol Biotechnol. 2017;33(11):204.

12. Tabor CW, Tabor H. Polyamines in microorganisms. Microbiol Rev. 1985;49(1):81–99.

13. Matsumoto M, Kibe R, Ooga T, Aiba Y, Kurihara S, Sawaki E, et al. Impact of Intestinal Microbiota on Intestinal Luminal Metabolome. Scientific Reports. 2012;2(1):233.

14. Milovic V. Polyamines in the gut lumen: bioavailability and biodistribution. Eur J Gastroenterol Hepatol. 2001;13(9):1021–5.

15. Nakamura A, Kurihara S, Takahashi D, Ohashi W, Nakamura Y, Kimura S, et al. Symbiotic polyamine metabolism regulates epithelial proliferation and macrophage differentiation in the colon. Nature Communications. 2021;12(1):2105.

16. Tofalo R, Cocchi S, Suzzi G. Polyamines and Gut Microbiota. Front Nutr. 2019;6:16.

17. Kitada Y, Muramatsu K, Toju H, Kibe R, Benno Y, Kurihara S, et al. Bioactive polyamine production by a novel hybrid system comprising multiple indigenous gut bacterial strategies. Sci Adv. 2018;4(6):eaat0062.

18. De Biase D, Pennacchietti E. Glutamate decarboxylase-dependent acid resistance in orally acquired bacteria: function, distribution and biomedical implications of the gadBC operon. Mol Microbiol. 2012;86(4):770–86.

19. Iyer R, Williams C, Miller C. Arginine-agmatine antiporter in extreme acid resistance in Escherichia coli. J Bacteriol. 2003;185(22):6556–61.

20. Chattopadhyay MK, Tabor H. Polyamines are critical for the induction of the glutamate decarboxylase-dependent acid resistance system in Escherichia coli. J Biol Chem. 2013;288(47):33559–70.

21. Otaru N, Ye K, Mujezinovic D, Berchtold L, Constancias F, Cornejo FA, et al. GABA Production by Human Intestinal Bacteroides spp.: Prevalence, Regulation, and Role in Acid Stress Tolerance. Front Microbiol. 2021;12:656895.

22. Sugiyama Y, Nara M, Sakanaka M, Gotoh A, Kitakata A, Okuda S, et al. Comprehensive analysis of polyamine transport and biosynthesis in the dominant human gut bacteria: Potential presence of novel polyamine metabolism and transport genes. Int J Biochem Cell Biol. 2017;93:52–61.

23. Lopez-Samano M, Beltran LFL, Sanchez-Thomas R, Davalos A, Villasenor T, Garcia-Garcia JD, et al. A novel way to synthesize pantothenate in bacteria involves beta-alanine synthase present in uracil degradation pathway. Microbiologyopen. 2020;9(4):e1006.

24. Hata J, Ohara T, Katakura Y, Shimizu K, Yamashita S, Yoshida D, et al. Association Between Serum beta-Alanine and Risk of Dementia. Am J Epidemiol. 2019;188(9):1637–45.

25. Ostfeld I, Ben-Zeev T, Zamir A, Levi C, Gepner Y, Springer S, et al. Role of beta-Alanine Supplementation on Cognitive Function, Mood, and Physical Function in Older Adults; Double-Blind Randomized Controlled Study. Nutrients. 2023;15(4).

26. Damiano MA, Bastianelli D, Al Dahouk S, Kohler S, Cloeckaert A, De Biase D, et al. Glutamate decarboxylase-dependent acid resistance in Brucella spp.: distribution and contribution to fitness under extremely acidic conditions. Appl Environ Microbiol. 2015;81(2):578–86.

27. McCormick DA. GABA as an inhibitory neurotransmitter in human cerebral cortex. J Neurophysiol. 1989;62(5):1018–27.

28. De Palma G, Shimbori C, Reed DE, Yu Y, Rabbia V, Lu J, et al. Histamine production by the gut microbiota induces visceral hyperalgesia through histamine 4 receptor signaling in mice. Sci Transl Med. 2022;14(655):eabj1895.

29. Del Rio B, Redruello B, Linares DM, Ladero V, Ruas-Madiedo P, Fernandez M, et al. The biogenic amines putrescine and cadaverine show in vitro cytotoxicity at concentrations that can be found in foods. Sci Rep. 2019;9(1):120.

30. Kovacs T, Miko E, Vida A, Sebo E, Toth J, Csonka T, et al. Cadaverine, a metabolite of the microbiome, reduces breast cancer aggressiveness through trace amino acid receptors. Sci Rep. 2019;9(1):1300.

31. Raiteri M, Del Carmine R, Bertollini A, Levi G. Effect of sympathomimetic amines on the synaptosomal transport of noradrenaline, dopamine and 5-hydroxytryptamine. Eur J Pharmacol. 1977;41(2):133–43.

32. Klein MO, Battagello DS, Cardoso AR, Hauser DN, Bittencourt JC, Correa RG. Dopamine: Functions, Signaling, and Association with Neurological Diseases. Cell Mol Neurobiol. 2019;39(1):31–59.

33. Berger M, Gray JA, Roth BL. The expanded biology of serotonin. Annu Rev Med. 2009;60:355–66.

34. Luqman A, Nega M, Nguyen MT, Ebner P, Gotz F. SadA-Expressing Staphylococci in the Human Gut Show Increased Cell Adherence and Internalization. Cell Rep. 2018;22(2):535–45.

35. Babusyte A, Kotthoff M, Fiedler J, Krautwurst D. Biogenic amines activate blood leukocytes via trace amine-associated receptors TAAR1 and TAAR2. J Leukoc Biol. 2013;93(3):387–94.

36. Xie Z, Miller GM. Beta-phenylethylamine alters monoamine transporter function via trace amine-associated receptor 1: implication for modulatory roles of trace amines in brain. J Pharmacol Exp Ther. 2008;325(2):617–28.

37. Mutuyemungu E, Singh M, Liu S, Rose DJ. Intestinal gas production by the gut microbiota: A review. Journal of Functional Foods. 2023;100:105367.

38. Momany C, Ghosh R, Hackert ML. Structural motifs for pyridoxal-5’-phosphate binding in decarboxylases: an analysis based on the crystal structure of the Lactobacillus 30a ornithine decarboxylase. Protein Sci. 1995;4(5):849–54.

39. Huynh QK, Snell EE. Pyruvoyl-dependent histidine decarboxylases. Preparation and amino acid sequences of the beta chains of histidine decarboxylase from Clostridium perfringens and Lactobacillus buchneri. J Biol Chem. 1985;260(5):2798–803.

40. Huynh QK, Snell EE. Pyruvoyl-dependent histidine decarboxylases. Comparative sequences of cysteinyl peptides of the enzymes from Lactobacillus 30a, Lactobacillus buchneri, and Clostridium perfringens. J Biol Chem. 1985;260(5):2794–7.

41. Recsei PA, Snell EE. Pyruvoyl-dependent histidine decarboxylases. Mechanism of cleavage of the proenzyme from Lactobacillus buchneri. J Biol Chem. 1985;260(5):2804–6.

42. Snell EE. Pyruvoyl-dependent histidine decarboxylase from Lactobacillus 30a: purification and properties. Methods Enzymol. 1986;122:128–35.

43. Nozaki S, Webb ME, Niki H. An activator for pyruvoyl-dependent l-aspartate alpha-decarboxylase is conserved in a small group of the gamma-proteobacteria including Escherichia coli. Microbiologyopen. 2012;1(3):298–310.

44. Tolbert WD, Graham DE, White RH, Ealick SE. Pyruvoyl-dependent arginine decarboxylase from Methanococcus jannaschii: crystal structures of the self-cleaved and S53A proenzyme forms. Structure. 2003;11(3):285–94.

45. Komori H, Nitta Y, Ueno H, Higuchi Y. Structural study reveals that Ser-354 determines substrate specificity on human histidine decarboxylase. J Biol Chem. 2012;287(34):29175–83.

46. Mou Z, Yang Y, Hall AB, Jiang X. The taxonomic distribution of histamine-secreting bacteria in the human gut microbiome. BMC Genomics. 2021;22(1):695.

47. Brown GM, Williamson JM. Biosynthesis of riboflavin, folic acid, thiamine, and pantothenic acid. Adv Enzymol Relat Areas Mol Biol. 1982;53:345–81.

48. Mahootchi E, Cannon Homaei S, Kleppe R, Winge I, Hegvik TA, Megias-Perez R, et al. GADL1 is a multifunctional decarboxylase with tissue-specific roles in beta-alanine and carnosine production. Sci Adv. 2020;6(29):eabb3713.

49. Tiedje KE, Stevens K, Barnes S, Weaver DF. Beta-alanine as a small molecule neurotransmitter. Neurochem Int. 2010;57(3):177–88.

50. Chen IA, Chu K, Palaniappan K, Ratner A, Huang J, Huntemann M, et al. The IMG/M data management and analysis system v.7: content updates and new features. Nucleic Acids Res. 2023;51(D1):D723–D32.

51. Forster SC, Kumar N, Anonye BO, Almeida A, Viciani E, Stares MD, et al. A human gut bacterial genome and culture collection for improved metagenomic analyses. Nat Biotechnol. 2019;37(2):186–92.

